# Accountable Prediction of Drug ADMET Properties with Molecular Descriptors

**DOI:** 10.1101/2022.06.29.115436

**Authors:** Nilavo Boral, Promita Ghosh, Adhish Goswami, Malay Bhattacharyya

**Author notes:** Correspondence to: Malay Bhattcharyya < >.

## Abstract

Drugs are chemical substances of low molecular weights that require to travel from the site of administration to the site of action. For safety and effective permeability, drugs are required to exhibit ideal absorption, distribution, metabolism, excretion and toxicity (ADMET) properties. Given the Simplified Molecular Input Line Entry System (SMILES) representation of drugs, we aim to predict their ADMET properties. By feeding molecular descriptors (as global features) to traditional machine learning models, we show that ADMET properties of drug molecules can be predicted with an accuracy competitive with the state-of-the-art deep learning models. We demonstrate that the proposed approach with only 31 molecular descriptors beats the state-of-the-art for 3 datasets. Moreover, it stands the second best for 2 other datasets, where none of the best provides a statistically significant improvement. We also demonstrate that two-dimensional descriptors can better represent absorption, distribution and excretion properties than the fingerprints widely used in the literature. However, they fail to distinguish metabolism and toxicity properties.

## 1. Introduction

A drug is a kind of chemical substance comprising small-molecules that causes changes in the physiology or psychology of an organism when consumed. Drugs are required to travel from the site of administration (e.g., oral, nasal, etc.) to the site of action (e.g., a tissue) (Arora et al., 2002).

Due to low molecular weights, drugs can comfortably travel within the body and pass across biological membrane at different levels. On completing the action, they decompose and depart the body. To perform these operations safely and efficiently, drugs are required to exhibit ideal absorption, distribution, metabolism, excretion and toxicity (ADMET) properties (Avdeef, 2012) (Dearden, 2007).

We aim to predict these ADMET properties of drug molecules with more explainability. Many types of fingerprints and descriptors (such as PubChem (Kim et al., 2020) fingerprints, extended connectivity fingerprints, Molecular ACCess System (MACCS) fingerprints (Polton, 1982), Mordred descriptors (Moriwaki et al., 2018) etc.) can be extracted and already used in previous ADMET prediction methods. In this paper, We show that only using 31 two-dimensional descriptors (Matter, 1997) as features, some ADMET related properties of drug can be predicted with high accuracy. Our prediction method is able to predict 3 properties (half life obach, vdss lombardo, clearance hepatocyte az) ranked first and 5 properties ranked in top 3 among current top models.

## 2. Problem Statement

The prediction of the ADMET properties plays an important role in the drug discovery and development process because these properties account for the failure of about 60% of all drugs in the clinical phases. Poor ADMET profile is the most prominent reason of failure in clinical trials. Thus, an early and accurate ADMET profiling during the discovery stage is a necessary condition for successful development of small-molecule candidate. In real-world discovery, the drug structures of interest evolve over time. Thus, ADMET prediction requires a model to generalize to a set of unseen drugs that are structurally distant to the known drug set.

## 3. Methods

Though there are a number of sophisticated deep learning approaches in existence, we have demonstrated that they miss including many important features. That is why, in this paper we propose, a simplistic approach of global features extraction method and some simple machine learning algorithms with optimised parameters to predict different ADMET properties of drug molecule.

### 3.1. Data Collection

In our study, the experimental values and corresponding basic information of drug molecule for ADMET properties prediction were collected from Therapeutics Data Commons (TDC) (Huang et al., 2021). We considered total 22 datasets for the current analysis (see Appendix A). The categorization of these datasets is highlighted in Fig. 1.

**Figure 1.**
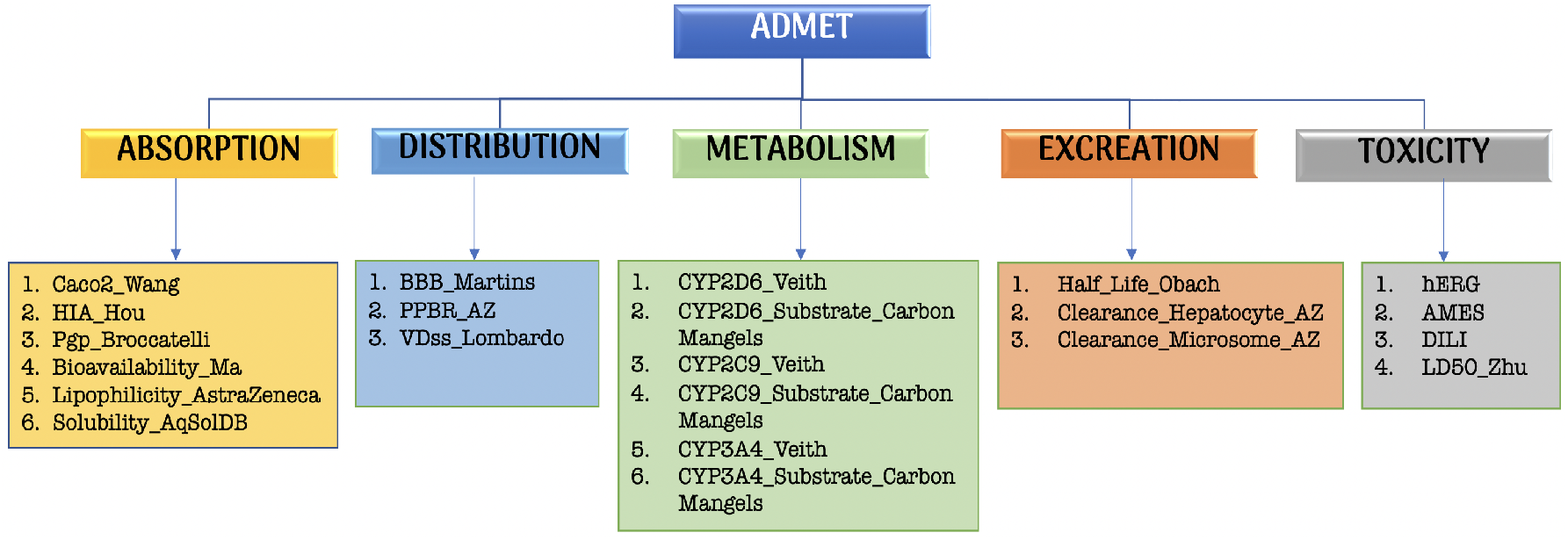
The 22 datasets related to different ADMET properties.

### 3.2. Dataset Split

For each dataset, TDC provides a separate test set (20%) and a training-validation set (80%).

#### Choosing the best model

We tried to employ different machine learning models (classifiation and regression tasks) to find out the best one. For this, we randomly split (to maintain the unbiasness/randomness of data at the time of model choosing) the training set of each dataset in training set (87.5%) and validation set (12.5%) using the TDC utility function. Then used training and validation set to find out the best predicting ML models for each dataset.

#### Testing on the independent test set

For a more rigorous analysis, we again split the training set (87.5%) and validation set (12.5%) using scaffold split (based on the scaffold of the molecules so that training/validation set is structurally different). We then trained the chosen model using both training and validation set, and finally test the model’s performance on the independent test set.

### 3.3. Feature Extraction

Drug SMILES (Simplified Molecular Input Line Entry System) (O’Boyle, 2012) strings were converted into molecular-data files (‘mol’ format) and fed into RDKit, an open-source cross-platform chemoinformatics toolkit. The tool has a built-in functionality for generating both compositional descriptors like MolWt, NumValenceElectrons, NumHDonor, etc. and topological molecular descriptors like FpDensity-Morgan1, FpDensityMorgan2, FpDensityMorgan3, etc. The molecular data files were then read by the Chem.Descriptors and Chem.Lipinski modules of RDKit to compute 31 molecular descriptors (see Appendix B) for each molecule in each dataset. This descriptors together can represent the overall behaviour of a drug.

Each feature can individually describe some behaviour of a whole drug molecule, so we can also call this global descriptors of a molecule. This global features improves the prediction of some ADMET properties of drug molecules.

### 3.4. Modeling Algorithm

In this study, different modeling algorithms were applied to develop regression or classification models for ADMET related properties: Linear Regression (for regression problem), Logistic Regression (for classification problem), KNN, Random Forest, Extra Tree, Bagging, AdaBoost, Decision Tree, Gradient Boosting, XGBoost.

- Linear Regression (Montgomery et al., 2021) shows the relationship between one dependent variable and one or more independent variable. How dependent variable changes with independent variables. Linear models generate a formula to create a best-fit line to predict unknown values.
- Logistic Regression (Kleinbaum et al., 2002) is a regression model. The model builds a regression model to predict the probability that a given data entry belongs to the category numbered as “1”. Just like Linear regression assumes that the data follows a linear function, Logistic regression models the data using the sigmoid function. Logistic regression becomes a classification technique only when a decision threshold is selected.
- The KNN (Guo et al., 2003) algorithm uses ‘feature similarity’ to predict the values of any new data points for both regression and classification problem. Which means that the new point is assigned a value based on how closely it resembles the points in the training set.
- Decision trees (Quinlan, 1987) is a non-parametric supervised learning method used for classification and regression. The goal is to create a model that predicts the value of a target variable by learning simple decision rules inferred from the data features.
- Bagging (Bühlmann & Yu, 2002) is an ensemble learning technique used in both regression and classification model. It is used to deal with bias-variance trade-offs and reduces the variance of a prediction model. It avoids overfitting of data.
- Random Forest (Breiman, 2001) is an ensemble of unpruned classification or regression trees created by using bootstrap samples of the training data and random feature selection in tree induction.
- Extra Tree (Bhati & Rai, 2020) is a type of ensemble learning technique that aggregates the results of different de-correlated decision trees. Extra Tree does not perform bootstrap aggregation like in the random forest. This model takes a random subset of data without replacement. Thus nodes are split on random splits and not on best splits.
- AdaBoost (Schapire, 2013) was the first really successful boosting algorithm. AdaBoost is short for Adaptive Boosting and is a very popular boosting technique that combines multiple “weak models” into a single “strong models”.
- Gradient Boosting (Natekin & Knoll, 2013) is a popular boosting algorithm. In gradient boosting, each predictor corrects its predecessor’s error. In contrast to Adaboost, the weights of the training instances are not tweaked, instead, each predictor is trained using the residual errors of predecessor as labels. There is a technique called the Gradient Boosted Trees whose base learner is CART (Classification and Regression Trees).
- XGBoost (Chen & Guestrin, 2016) is an implementation of Gradient Boosted decision trees. In this algorithm, decision trees are created in sequential form. Weights play an important role in XGBoost. Weights are assigned to all the independent variables which are then fed into the decision tree which predicts results. The weight of variables predicted wrong by the tree is increased and these variables are then fed to the second decision tree. These individual classifiers/predictors then ensemble to give a strong and more precise model.

We trained these models with default parameters on training set and then we compared statistical performances (MAE for Regression datasets, Accuracy for Classification datasets) of all models on validation set to choose top 2 best models, then using RandomSearchCV we optimised parameters of that top 2 ML models to find out a best model for each ADMET property prediction problem.

### 3.5. Performance Analysis

Further study was performed to verify our method’s robustness and predictive ensure that the derived model from the training set has good ability. For generalization ability, we fitted our model on training and validation set and finally checked the statistical performance of testing set. We did this five times to calculate average and standard deviation of prediction results.

## 4. Results

Some of the datasets used for predicting ADMET properties are posed as binary classification problems and the rest as regression problems. We choose AUROC (area under ROC curve) and AUPRC (area under PRC curve) as the performance metrics for the binary classification problems (Lei et al., 2019), and Mean Absolute Error (MAE) (Chai & Draxler, 2014) and Spearman’s correlation coefficient for regression problems. We apply all the machine learning models mentioned earlier. Average and standard deviation of their performances across five independent runs on the test sets are reported in Tables 1-5. The results are compared with all of the state-of-the-art methods reporting their performance as a part of the ADMET Benchmark Group challenge^1^. We used The Google Colaboratory^2^ with default runtime (without any virtual hardware accelerator like GPU or TPU) for all our experiments. For data collection as well as data set splitting, we employed utility functions from TDC.

**Table 1.**
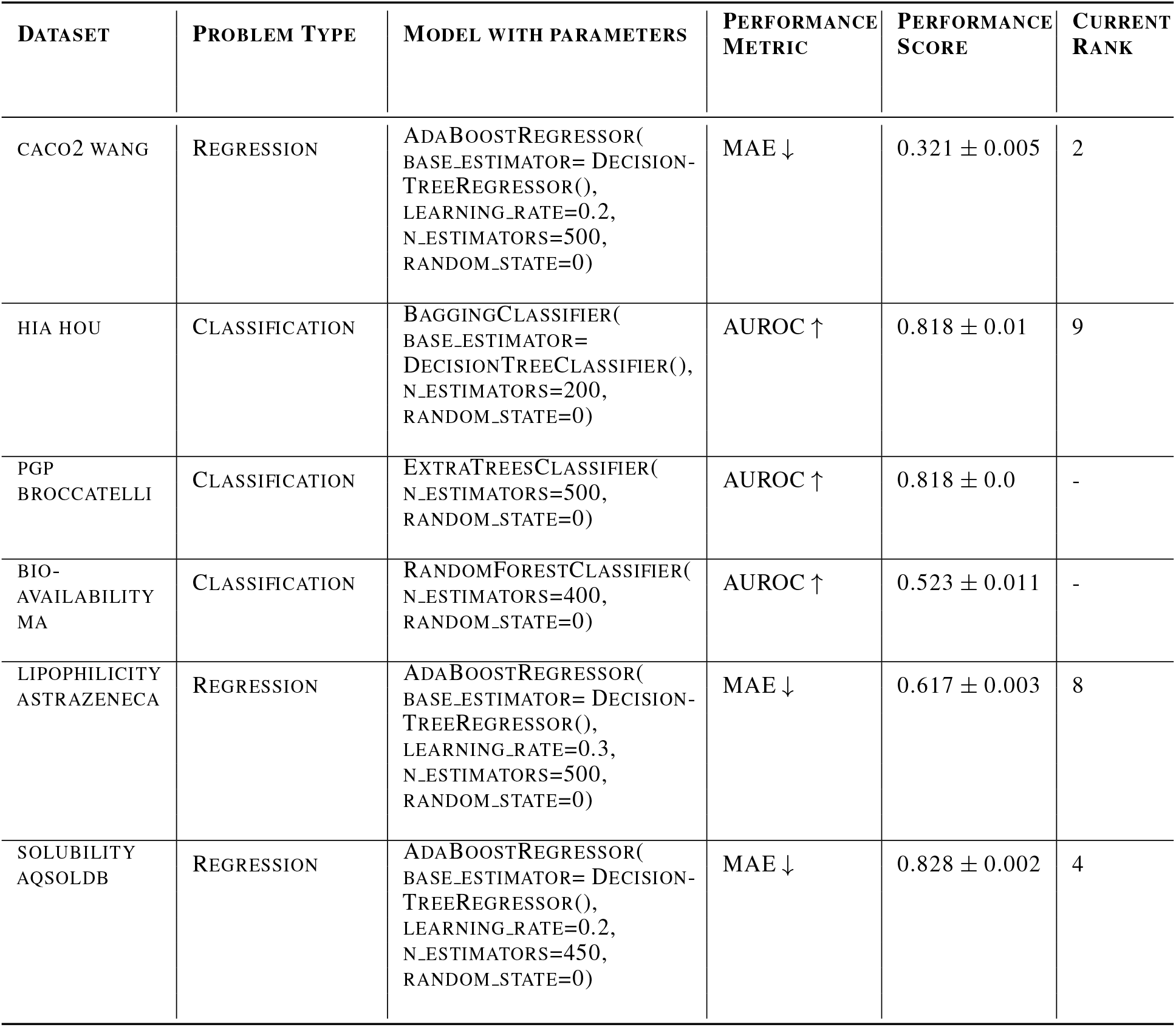
Comparative performance of predicting ABSORPTION related property in the ADMET Benchmark Group challenge. Arrows (*↑, ↓*) in performance metric indicate the direction of superiority.

### 4.1. Results for Absorption related drug properties

There are 3 regression and 3 classification problems defined around the 6 datasets related to Absorption properties. We observe that for all the regression problems adaboost regressor performs the best. On the other side, for the classification problems, we observe that there is no such single model which works the best across all the datasets. On comparing the chosen best models with the state-of-the-art, we observe that our method is able to rank within top 10 in the ADMET Benchmark Group challenge for several datasets. To be precise, it ranks within top 10 for all the three regression datasets that includes a second best case. However, for only one classification dataset it performs well (see Table 1).

### 4.2. Results for Distribution related drug properties

There are 2 regression and 1 classification problems defined around the 3 datasets related to Distribution Properties. Comparing our best model with the state-of-the-art models reported in the ADMET Benchmark Group challenge, we observe that for both the regression problems ranks are obtained within the top 2, including one case depicting the rank 1. But for the only classification problem, our method holds a rank of 10 (see Table 2).

**Table 2.** Comparative performance of predicting DISTRIBUTION related property in the ADMET Benchmark Group challenge. Arrows (*↑, ↓*) in performance metric indicate the direction of superiority.

| DATASET | PROBLEM TYPE | MODEL WITH PARAMETERS | PERFORMANCE METRIC | PERFORMANCE SCORE | CURRENT RANK |
| --- | --- | --- | --- | --- | --- |
| BBB MARTINS | CLASSIFICATION | BAGGINGCLASSIFIER(<br>BASE_ESTIMATOR=<br>DECISIONTREECLASSIFIER(),<br>N_ESTIMATORS=100,<br>RANDOM_STATE=0) | AUROC ↑ | $0.811 \pm 0.013$ | 10 |
| PPBR AZ | REGRESSION | GRADIENTBOOSTING-REGRESSOR(<br>LEARNING_RATE=0.005,<br>N_ESTIMATORS=500,<br>RANDOM_STATE=0) | MAE ↓ | $9.185 \pm 0.0$ | 2 |
| VDSS<br>LOMBARDO | REGRESSION | ADABOOSTREGRESSOR(<br>BASE_ESTIMATOR= DECISION-TREEREGRESSOR(),<br>LEARNING_RATE=0.005,<br>N_ESTIMATORS=200,<br>RANDOM_STATE=0) | SPEARMAN ↑ | $0.627 \pm 0.01$ | 1 |

### 4.3. Results for Metabolism related drug properties

There are 6 datasets in metabolism related properties and all of them are classification problems. For all these datasets ensemble learning techniques are found to perform well among all the models we employed. On comparing this with all available top models in the ADMET Benchmark Group challenge, we observe that these models hardly manage to get a rank within top 10, except for one case (see Table 3).

**Table 3.** Comparative performance of predicting METABOLISM related property in the ADMET Benchmark Group challenge. Arrows (*↑, ↓*) in performance metric indicate the direction of superiority.

| DATASET | PROBLEM TYPE | MODEL WITH PARAMETERS | PERFORMANCE METRIC | PERFORMANCE SCORE | CURRENT RANK |
| --- | --- | --- | --- | --- | --- |
| CYP2D6 VEITH | CLASSIFICATION | EXTRATREESCLASSIFIER(<br>N_ESTIMATORS=250,<br>RANDOM_STATE=0) | AUPRC ↑ | $0.358 \pm 0.0$ | - |
| CYP3A4 VEITH | CLASSIFICATION | XGBCLASSIFIER(<br>N_ESTIMATORS=400) | AUPRC ↑ | $0.654 \pm 0.0$ | - |
| CYP2C9 VEITH | CLASSIFICATION | BAGGINGCLASSIFIER(<br>BASE_ESTIMATOR=<br>DECISIONTREECLASSIFIER(),<br>N_ESTIMATORS=350,<br>RANDOM_STATE=0) | AUPRC ↑ | $0.556 \pm 0.0$ | - |
| CYP2D6<br>SUBSTRATE<br>CARBONMANGELS | CLASSIFICATION | BAGGINGCLASSIFIER(<br>BASE_ESTIMATOR=<br>DECISIONTREECLASSIFIER(),<br>N_ESTIMATORS=250,<br>RANDOM_STATE=0) | AUPRC ↑ | $0.478 \pm 0.018$ | - |
| CYP3A4<br>SUBSTRATE<br>CARBONMANGELS | CLASSIFICATION | XGBCLASSIFIER(<br>LEARNING_RATE=0.005,<br>N_ESTIMATORS=150) | AUROC ↑ | $0.605 \pm 0.0$ | 7 |
| CYP2C9<br>SUBSTRATE<br>CARBONMANGELS | CLASSIFICATION | BAGGINGCLASSIFIER(<br>BASE_ESTIMATOR=<br>DECISIONTREECLASSIFIER(),<br>N_ESTIMATORS=500,<br>RANDOM_STATE=0) | AUPRC ↑ | $0.281 \pm 0.0$ | - |

### 4.4. Results for Excretion related drug properties

There are 3 regression datasets related to Excretion properties. We observe that tree based models, specifically Adaboost and extra tree regression models, work well for these datasets. On comparing our best performing models with the state-of-the-art results reported in the ADMET Benchmark Group challenge, we observe that traditional machine learning models perform significantly well for all the regression datasets (see Table 4). This possibly highlights that two-dimensional molecular descriptors are effective features for characterizing excretion property of drug molecules.

**Table 4.** Comparative performance of predicting EXCRETION related property in the ADMET Benchmark Group challenge. Arrows (*↑, ↓*) in performance metric indicate the direction of superiority.

| DATASET | PROBLEM TYPE | MODEL WITH PARAMETERS | PERFORMANCE METRIC | PERFORMANCE SCORE | CURRENT RANK |
| --- | --- | --- | --- | --- | --- |
| HALF LIFE<br>OBACH | REGRESSION | ADABOOSTREGRESSOR(<br>BASE_ESTIMATOR= DECISION-TREEREGRESSOR(),<br>LEARNING_RATE=0.08,<br>N_ESTIMATORS=350,<br>RANDOM_STATE=0) | SPEARMAN ↑ | $0.438 \pm 0.011$ | 1 |
| CLEARANCE<br>MICROSOME<br>AZ | REGRESSION | ADABOOSTREGRESSOR(<br>BASE_ESTIMATOR= DECISION-TREEREGRESSOR(),<br>LEARNING_RATE=0.3,<br>N_ESTIMATORS=350,<br>RANDOM_STATE=0) | SPEARMAN ↑ | $0.518 \pm 0.005$ | 8 |
| CLEARANCE<br>HEPATOCYTE<br>AZ | REGRESSION | EXTRATREESREGRESSOR(<br>N_ESTIMATORS=500,<br>RANDOM_STATE=0) | SPEARMAN ↑ | $0.44 \pm 0.003$ | 1 |

### 4.5. Results for Toxicity related drug properties

There are 1 regression and 3 classification problems defined around the 4 datasets related to Toxicity properties. On applying the commonly used machine learning models mentioned before, we observe that none of the chosen machine learning models exhibit a consistent performance for the classification datasets. However, for the regression tasks, our models manage to get a rank 4 among the present top models included in the ADMET Benchmark Group challenge (see Table 5).

**Table 5.** Comparative performance of predicting TOXICITY related property in the ADMET Benchmark Group challenge. Arrows (*↑, ↓*) in performance metric indicate the direction of superiority.

| DATASET | PROBLEM TYPE | MODEL WITH PARAMETERS | PERFORMANCE METRIC | PERFORMANCE SCORE | CURRENT RANK |
| --- | --- | --- | --- | --- | --- |
| HERG | CLASSIFICATION | BAGGINGCLASSIFIER(<br>BASE_ESTIMATOR=<br>DECISIONTREECLASSIFIER(),<br>N_ESTIMATORS=150,<br>RANDOM_STATE=0) | AUROC ↑ | $0.715 \pm 0.011$ | - |
| AMES | CLASSIFICATION | EXTRA TREES CLASSIFIER(<br>N_ESTIMATORS=500,<br>RANDOM_STATE=0) | AUROC ↑ | $0.716 \pm 0.0$ | - |
| DILI | CLASSIFICATION | KNEIGHBORSCLASSIFIER(<br>N_NEIGHBORS=4) | AUROC ↑ | $0.7 \pm 0.0$ | - |
| LD50 ZHU | REGRESSION | EXTRA TREES REGRESSOR(<br>N_ESTIMATORS=350,<br>RANDOM_STATE=0) | MAE ↓ | $0.636 \pm 0.001$ | 4 |

To better understand the importance of the features across all the datasets, we have utilized the internal feature importance ranking function in Extra-Tree model. All features were applied and default parameters were used to build the model. The results are reported in Table 6.

**Table 6.**
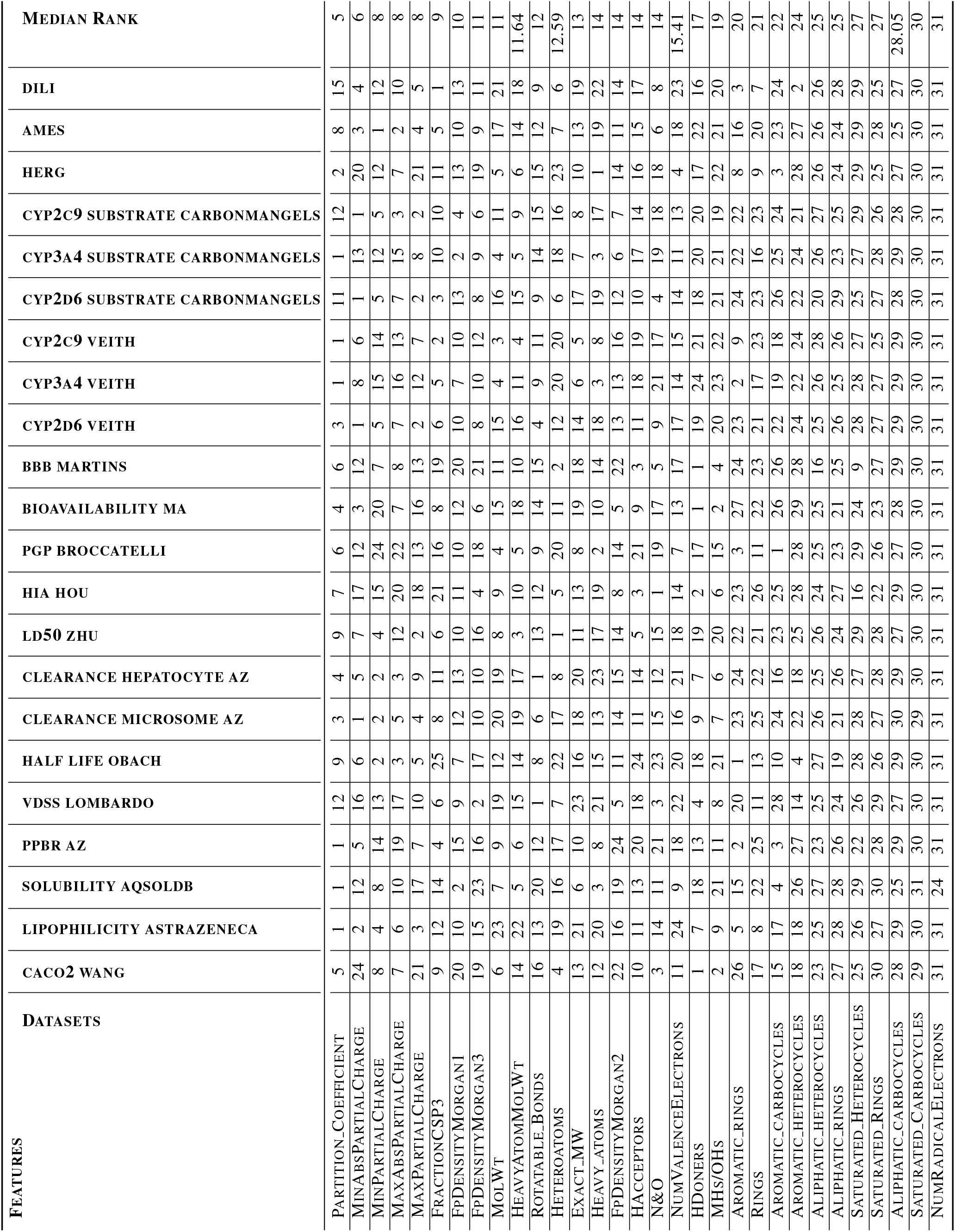
Ranking of descriptors (features) based on their prediction capability of ADMET properties across all the datasets. The features are ordered by their average rank.

## 5. Discussion

A significant of the state-of-the-art models (taking part in the ADMET Benchmark Group challenge) that we compared our results with are based on fingerprint based features. The limited rest of them uses a large amount of molecular descriptors, thereby giving no clue about what kind of descriptors might be appropriate for predicting what type of ADMET property. A deeper look at the prediction results across all of the ADMET properties (Tables 1-5) reveal that two-dimensional descriptors can better represent the properties like absorption, distribution and excretion. Note that the descriptors that are used in the current analysis are based on oral bioavailability (Veber et al., 2002) of drug molecules. This might be the reason of their success for the said three properties. On the other hand, metabolism is related to biochemical properties of the drug molecules and their clinical relevance is difficult to assess from the molecular descriptors only. Moreover, the toxicity property, being connected with the drug target interactions (Yamanishi et al., 2010), cannot be characterized suitably through those descriptors. We have seen that the traditional machine learning models chosen by us are more effective in solving regression problems than classification because of using molecular descriptors. This is possibly because most of the regression problems are posed from datasets pertaining to absorption, distribution and excretion properties.

It appears from the median ranking of the descriptors (see Table 6) across all the experiments that some Lipinski parameters are crucial in reflecting the ADMET properties. This includes octanol-water partition coefficient (PARTITION COEFFICIENT), fraction of C atoms that are SP3 hybridized (FRACTIONCSP3), partial charge (both minimum and maximum), and topological molecular descriptors (like FPDENSITYMORGAN1 and 3). A recent analysis has shown that most of these are two-dimensional topological/topochemical properties that provide useful information about the molecular surface and its potential interactions with the binding species (Chaube et al., 2020). Interestingly, topological molecular descriptors generate similarity fingerprints using certain chemical and connectivity attributes of atoms (Riniker & Landrum, 2013). Hence, they provide a better accountability in predicting the ADMET properties of drugs.

## 6. Conclusion

Through this study we worked only with 31 two-dimensional global molecular descriptors. For that, our method is very less expensive in terms of time and computational power. These 31 features with some traditional machine learning models are sufficient to predict 3 ADMET properties more accurately than any of the current existing methods or models. Though we focus only on twodimensional descriptors in this work, higher dimensional descriptors are also worthy to look into in future.

## Availability

All the codes related to our work (including data import, features extraction, feature selection, modeling and performance checking) are publicly available from the GitHub link: https://github.com/NilavoBoral/Therapeutics-Data-Commons.

## Appendix A

The 22 datasets that we considered from TDC for the current analysis are described hereunder (Huang et al., 2021).

- **Datasets Related to Absorption Property ():**
  1. **caco2-wang:** The human colon epithelial cancer cell line, Caco-2, is used as an in vitro model to simulate the human intestinal tissue. The experimental result on the rate of drug passing through the Caco-2 cells can approximate the rate at which the drug permeates through the human intestinal tissue.
  2. **hia-hou:** When a drug is orally administered, it needs to be absorbed from the human gastrointestinal system into the bloodstream of the human body. This ability of absorption is called human intestinal absorption (HIA) and it is crucial for a drug to be delivered to the target.
  3. **pgp-broccatelli:** P-glycoprotein (Pgp) is an ABC transporter protein involved in intestinal absorption, drug metabolism, and brain penetration, and its inhibition can seriously alter a drug’s bioavailability and safety. In addition, inhibitors of Pgp can be used to overcome multi-drug resistance.
  4. **bioavailability-ma:** Oral bioavailability is defined as the rate and extent to which the active ingredient or active moiety is absorbed from a drug product and becomes available at the site of action.
  5. **lipophilicity-astrazeneca:** Lipophilicity measures the ability of a drug to dissolve in a lipid (e.g. fats, oils) environment. High lipophilicity often leads to high rate of metabolism, poor solubility, high turn-over, and low absorption.
  6. **solubility-aqsoldb:** Aqeuous solubility measures a drug’s ability to dissolve in water. Poor water solubility could lead to slow drug absorptions, inadequate bioavailablity and even induce toxicity. More than 40% of new chemical entities are not soluble.
- **Datasets Related to Distribution Property:**
  1. **bbb-martins:** As a membrane separating circulating blood and brain extracellular fluid, the blood-brain barrier (BBB) is the protection layer that blocks most foreign drugs. Thus the ability of a drug to penetrate the barrier to deliver to the site of action forms a crucial challenge in development of drugs for central nervous system.
  2. **ppbr-az:** The human plasma protein binding rate (PPBR) is expressed as the percentage of a drug bound to plasma proteins in the blood. This rate strongly affect a drug’s efficiency of delivery. The less bound a drug is, the more efficiently it can traverse and diffuse to the site of actions.
  3. **vdss-lombardo:** The volume of distribution at steady state (VDss) measures the degree of a drug’s concentration in body tissue compared to concentration in blood. Higher VD indicates a higher distribution in the tissue and usually indicates the drug with high lipid solubility, low plasma protein binidng rate.
- **Datasets Related to Metabolism Property:**
  1. **cyp2d6-veith:** The CYP P450 genes are involved in the formation and breakdown (metabolism) of various molecules and chemicals within cells. Specifically, CYP2D6 is primarily expressed in the liver. It is also highly expressed in areas of the central nervous system, including the substantia nigra.
  2. **cyp3a4-veith:** The CYP P450 genes are involved in the formation and breakdown (metabolism) of various molecules and chemicals within cells. Specifically, CYP3A4 is an important enzyme in the body, mainly found in the liver and in the intestine. It oxidizes small foreign organic molecules (xenobiotics), such as toxins or drugs, so that they can be removed from the body.
  3. **cyp2c9-veith:** The CYP P450 genes are involved in the formation and breakdown (metabolism) of various molecules and chemicals within cells. Specifically, the CYP P450 2C9 plays a major role in the oxidation of both xenobiotic and endogenous compounds.
  4. **cyp2d6-substrate-carbonmangels:** CYP2D6 is primarily expressed in the liver. It is also highly expressed in areas of the central nervous system, including the substantia nigra.
  5. **cyp3a4-substrate-carbonmangels:** CYP3A4 is an important enzyme in the body, mainly found in the liver and in the intestine. It oxidizes small foreign organic molecules (xenobiotics), such as toxins or drugs, so that they can be removed from the body.
  6. **cyp2c9-substrate-carbonmangels:** CYP P450 2C9 plays a major role in the oxidation of both xenobiotic and endogenous compounds. Substrates are drugs that are metabolized by the enzyme.
- **Datasets Related to Excretion Properties:**
  1. **half-life-obach:** Half life of a drug is the duration for the concentration of the drug in the body to be reduced by half. It measures the duration of actions of a drug.
  2. **clearance-hepatocyte-az:** Hepatocytes, the major parenchymal cells in the liver, play pivotal roles in drug metabolism, detoxification and excretion. They also activate innate immunity against invading microorganisms by secreting innate immunity proteins that directly kills bacteria and block iron uptake by bacteria. They determine the in vitro intrinsic clearance of any drug compound.
  3. **clearance-microsome-az:** Located in the liver, Microsomes are subcellular fractions, that contains membrane bound drug metabolishing enzymes, can determine the in-vitro intrinsic clearance of a drug molecule. Microsomes are pooled from multiple donors to minimise the effect of inter-individual variability. Liver microsomes are more predictive of in vivo clearance than hepatocytes, when in vitro intrinsic clearance in microsomes is faster than hepatocytes.
- **Datasets Related to Toxicity Properties:**
  1. **herg:** Human ether-a’-go-go related gene (hERG) is crucial for the coordination of the heart’s beating. Thus, if a drug blocks the hERG, it could lead to severe adverse effects. Therefore, reliable prediction of hERG liability in the early stages of drug design is quite important to reduce the risk of cardiotoxicity-related attritions in the later development stages.
  2. **ames:** Mutagenicity means the ability of a drug to induce genetic alterations. Drugs that can cause damage to the DNA can result in cell death or other severe adverse effects. Nowadays, the most widely used assay for testing the mutagenicity of compounds is the Ames experiment which was invented by a professor named Ames. The Ames test is a short-term bacterial reverse mutation assay detecting a large number of compounds which can induce genetic damage and frameshift mutations.
  3. **dili:** Drug-induced liver injury (DILI) is fatal liver disease caused by drugs and it has been the single most frequent cause of safety-related drug marketing withdrawals for the past 50 years (e.g. iproniazid, ticrynafen, benoxaprofen).
  4. **acute-toxicity-LD50:** Acute toxicity LD50 measures the most conservative dose that can lead to lethal adverse effects. The higher the dose, the more lethal of a drug.

## Appendix B

The 31 descriptors (used as features) are listed below.

- Exact molecular weight,
- FpDensityMorgan1,
- FpDensityMorgan2,
- FpDensityMorgan3,
- Average molecular weight of the molecule ignoring hydrogens,
- Average molecular weight,
- Number of radical electrons,
- Number of valence electrons,
- Partition coefficient,
- Fraction of C atoms that are SP3 hybridized,
- Number of heavy atoms,
- Number of NHs or OHs,
- Number of Nitrogens and Oxygens,
- Number of aliphatic carbocycles,
- Number of aliphatic heterocycles,
- Number of aliphatic rings,
- Number of aromatic carbocycles,
- Number of aromatic heterocycles,
- Number of aromatic rings,
- Number of Hydrogen Bond Acceptors,
- Number of Hydrogen Bond Donors,
- Number of Heteroatoms,
- Number of Rotatable Bonds,
- Number of saturated carbocycles,
- Number of saturated heterocycles,
- Number of saturated rings,
- Number of rings.

## Footnotes

1 https://tdcommons.ai/benchmark/admet_group/overview

2 https://colab.research.google.com

## References

Arora, P., Sharma, S., and Garg, S. Permeability issues in nasal drug delivery. Drug discovery today, 7(18):967–975,. 2002.

Avdeef, A. Absorption and drug development: solubility, permeability, and charge state. John Wiley & Sons, 2012.

Bhati, B. S. and Rai, C. Ensemble based approach for intrusion detection using extra tree classifier. In Intelligent computing in engineering, pp. 213–220. Springer, 2020.

Breiman, L. Random forests. Machine learning, 45(1): 5–32, 2001.

Bühlmann, P. and Yu, B. Analyzing bagging. The annals of Statistics, 30(4):927–961, 2002.

Chai, T. and Draxler, R. R. Root mean square error (rmse) or mean absolute error (mae). Geoscientific Model Development Discussions, 7(1):1525–1534, 2014.

Chaube, S., Goverapet Srinivasan, S., and Rai, B. Applied machine learning for predicting the lanthanide-ligand binding affinities. Scientific reports, 10(1):1–11, 2020.

Chen, T. and Guestrin, C. Xgboost: A scalable tree boosting system. In Proceedings of the 22nd acm sigkdd international conference on knowledge discovery and data mining, pp. 785–794, 2016.

Dearden, J. C. In silico prediction of admet properties: how far have we come? Expert Opinion on Drug Metabolism & Toxicology, 3(5):635–639, 2007.

Guo, G., Wang, H., Bell, D., Bi, Y., and Greer, K. Knn model-based approach in classification. In OTM Con-federated International Conferences” On the Move to Meaningful Internet Systems”, pp. 986–996. Springer, 2003.

Huang, K., Fu, T., Gao, W., Zhao, Y., Roohani, Y., Leskovec, J., Coley, C. W., Xiao, C., Sun, J., and Zitnik, M. Therapeutics data commons: Machine learning datasets and tasks for drug discovery and development. arXiv preprint 2102.09548, 2021.

Kim, S., Chen, J., Cheng, T., Gindulyte, A., He, J., He, S., Li, Q., Shoemaker, B. A., Thiessen, P. A., Yu, B., Zaslavsky, L., Zhang, J., and Bolton, E. E. PubChem in 2021: new data content and improved web interfaces. Nucleic Acids Research, 49(D1):D1388–D1395, 11 2020. ISSN 0305-1048. doi: 10.1093/nar/gkaa971. URL https://doi.org/10.1093/nar/gkaa971.

Kleinbaum, D. G., Dietz, K., Gail, M., Klein, M., and Klein, M. Logistic regression. Springer, 2002.

Lei, V. J., Kennedy, E. H., Luong, T., Chen, X., Polsky, D. E., Volpp, K. G., Neuman, M. D., Holmes, J. H., Fleisher, L. A., and Navathe, A. S. Model performance metrics in assessing the value of adding intraoperative data for death prediction: Applications to noncardiac surgery. In MedInfo, pp. 223–227, 2019.

Matter, H. Selecting optimally diverse compounds from structure databases: a validation study of two-dimensional and three-dimensional molecular descriptors. Journal of Medicinal Chemistry, 40(8):1219–1229, 1997.

Montgomery, D. C., Peck, E. A., and Vining, G. G. Introduction to linear regression analysis. John Wiley and Sons, 2021.

Moriwaki, H., Tian, Y.-S., Kawashita, N., and Takagi, T. Mordred: a molecular descriptor calculator. Journal of cheminformatics, 10(1):1–14, 2018.

Natekin, A. and Knoll, A. Gradient boosting machines, a tutorial. Frontiers in neurorobotics, 7:21, 2013.

O’Boyle, N. M. Towards a universal smiles representation-a standard method to generate canonical smiles based on the inchi. Journal of cheminformatics, 4(1):1–14, 2012.

Polton, D. Installation and operational experiences with maccs (molecular access system). Online Review, 1982.

Quinlan, J. R. Simplifying decision trees. International journal of man-machine studies, 27(3):221–234, 1987.

Riniker, S. and Landrum, G. A. Similarity maps-a visualization strategy for molecular fingerprints and machinelearning methods. Journal of cheminformatics, 5(1):1–7, 2013.

Schapire, R. E. Explaining adaboost. In Empirical inference, pp. 37–52. Springer, 2013.

Veber, D. F., Johnson, S. R., Cheng, H.-Y., Smith, B. R., Ward, K. W., and Kopple, K. D. Molecular properties that influence the oral bioavailability of drug candidates. Journal of medicinal chemistry, 45(12):2615–2623, 2002.

Yamanishi, Y., Kotera, M., Kanehisa, M., and Goto, S. Drugtarget interaction prediction from chemical, genomic and pharmacological data in an integrated framework. Bioinformatics, 26(12):i246–i254, 2010.

